# Supraclavicular brown adipocytes originate from *Tbx1*^+^ myoprogenitors

**DOI:** 10.1101/2023.06.16.545229

**Authors:** Zan Huang, Chenxin Gu, Zengdi Zhang, Rini Arianti, Aneesh Swaminathan, Kevin Tran, Alex Battist, Endre Kristóf, Hai-Bin Ruan

**Affiliations:** Laboratory of Gastrointestinal Microbiology, Jiangsu Key Laboratory of Gastrointestinal Nutrition and Animal Health, College of Animal Science and Technology, Nanjing Agricultural University, Nanjing, Jiangsu 210095, China; National Center for International Research on Animal Gut Nutrition, Nanjing Agricultural University, Nanjing, Jiangsu 210095, China; Department of Integrative Biology and Physiology, University of Minnesota Medical School, Minneapolis, MN 55455, USA; Department of Biochemistry and Molecular Biology, Faculty of Medicine, University of Debrecen, Debrecen 4024, Hungary; Center for Immunology, University of Minnesota Medical School, Minneapolis, MN 55455, USA

**Author notes:** Correspondence: H.-B. R.

## Abstract

Brown adipose tissue (BAT) dissipates energy as heat, contributing to temperature control, energy expenditure, and systemic homeostasis. In adult humans, BAT mainly exists in supraclavicular areas and its prevalence is associated with cardiometabolic health. However, the developmental origin of supraclavicular BAT remains unknown. Here, using genetic fate mapping in mice, we demonstrate that supraclavicular brown adipocytes do not develop from the *Pax3*^+^/*Myf5*^+^ epaxial dermomyotome that gives rises to interscapular BAT. Instead, the *Tbx1^+^* lineage that specifies the pharyngeal mesoderm marks the majority of supraclavicular brown adipocytes. *Tbx1^Cre^*-mediated ablation of peroxisome proliferator-activated receptor gamma (PPARγ) or PR/SET Domain 16 (PRDM16), components of the transcriptional complex for brown fat determination, leads to supraclavicular BAT paucity or dysfunction, thus rendering mice more sensitive to cold exposure. Moreover, human deep neck BAT expresses higher levels of the *TBX1* gene than subcutaneous neck white adipocytes. Taken together, our observations reveal location-specific developmental origins of BAT depots and call attention to *Tbx1^+^*lineage cells when investigating human relevant supraclavicular BAT.

## INTRODUCTION

Brown adipose tissue (BAT) is a thermogenic organ found in almost all mammals that dissipates energy as heat, thus contributing to homeostatic regulation of body temperature and metabolic physiology. In newborn humans, the predominant BAT depot is located in the interscapular region (iBAT). Through poorly understood mechanisms (Huang et al., 2022), iBAT undergoes progressive involution, scatters around the back during adolescence, and becomes undetectable in most adults (Heaton, 1972; Ruan, 2020; Sidossis and Kajimura, 2015). In adult humans, metabolically active BAT instead exists in cervical and supraclavicular areas, collectively referred as neck BAT (Cypess et al., 2009; Saito et al., 2009; van Marken Lichtenbelt et al., 2009; Virtanen et al., 2009; Zingaretti et al., 2009). The prevalence of neck BAT is inversely associated with body mass index (Betz and Enerback, 2011; Tam et al., 2012) and declines as a function of age (Saito *et al*., 2009; van Marken Lichtenbelt and Schrauwen, 2011; Yoneshiro et al., 2011), indicating the potential involvement of neck BAT dysfunction in the development of obesity and related metabolic disorders (Becher et al., 2021). However, the lineage origins and mechanisms for age-dependent functional decline of neck BAT remain almost unknown.

In the past decade, tremendous efforts have been made in our understanding of the development, recruitment, and activation of BAT. Nonetheless, most mechanistic studies were performed on rodent iBAT, due to its large size and easy accessibility. BAT and skeletal muscle have shared metabolic features and embryonic origins. Genetic fate mapping experiments in mice demonstrate that the dermomyotome regions of the somites, marked by the expression of transcription factors including *Pax3*, *Pax7*, *Meox1* and *Myf5*, gives rises to most fat cells within the interscapular and retroperitoneal adipose depots (Liu et al., 2013; Sanchez-Gurmaches and Guertin, 2014; Sanchez-Gurmaches et al., 2012; Seale et al., 2008; Sebo et al., 2018; Shan et al., 2013). The fact that these lineages trace to dorsal-anterior-located muscle, brown and white adipocytes suggests that they are location markers, rather than identity markers. Therefore, it is unlikely, although not tested or reported, that *Pax3^+^*/*Myf5*^+^ myoprogenitors form brown adipocytes in ventral neck BAT that has a very distinct location compared to dorsal-anterior BAT.

In vertebrates, head and neck muscles arise from the unsegmented cranial mesoderm, in distinction to somite-derived trunk muscles (Yahya et al., 2020). Transcriptional factors such as *Tbx1*, *Ptx2*, and *Islet1* specify the cardiopharyngeal mesoderm (CPM) that gives rise to muscles of the head and heart (Grimaldi et al., 2022; Heude et al., 2018; Lescroart et al., 2010). Supraclavicular BAT (scBAT) in mice is located in a region analogous to human neck BAT (**Figure 1A**). Though smaller than iBAT, subscapular and supraspinal (also termed as posterior cervical) BAT depots in the dorsal trunk (**Figure 1B**), scBAT possesses similar thermogenic activity and regulation (Mo et al., 2017; Shi et al., 2021). However, the developmental origins of scBAT adipocytes have not been defined. We hypothesized that the CPM also contributes to connective tissues in the neck region, including scBAT. In this study, taking advantage *Pax3^Cre^*, *Myf5^Cre^*, and *Tbx1^Cre^* -mediated lineage tracing and gene ablation, we identified the location-specific myogenic progenitors for scBAT versus iBAT in mice. Importantly, scBAT identity as *Tbx1*-progeny appears to be true in humans as well. This knowledge can be leverage in the future to investigate location-dependent functions of BAT and to target scBAT specifically for metabolic improvements.

**Figure 1.**
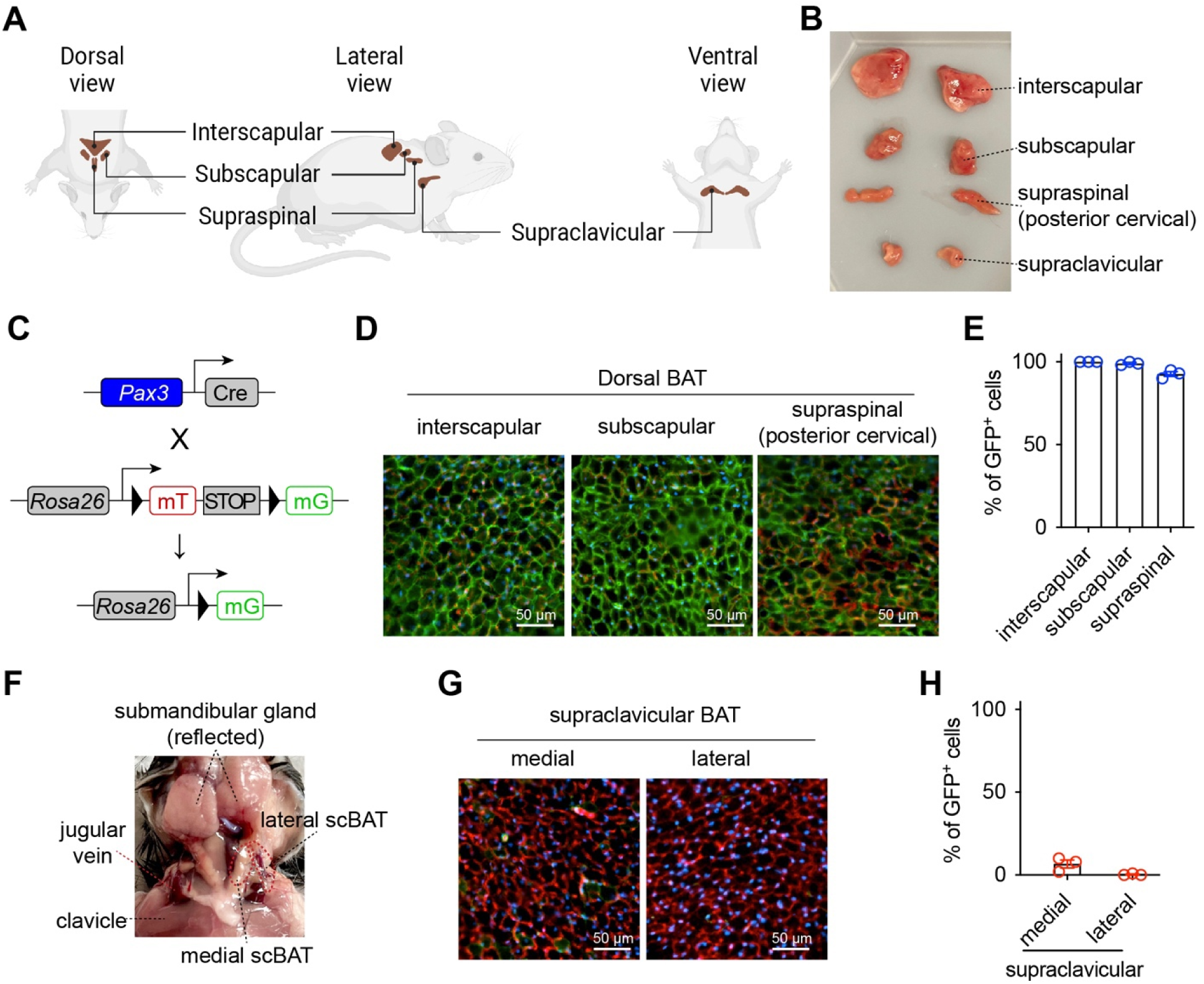
scBAT does not arise from *Pax3*^+^ progenitor cells. **(A)** Schematic representation of the location of peri-scapular and neck BAT in mice. **(B)** Representative photo of major BAT depots examined in this study. **(C)** Generation of the reporter mice for *Pax3*^+^ cells. **(D)** Fluorescent images of dorsal BAT depots from 5-week-old male *Pax3-mTmG* mice (green = mG, red = mT, blue = DAPI, Scale = 50 μm). **(E)** Frequency of GFP^+^ cells within each BAT depot (n = 3). **(F)** Anatomic location of supraclavicular BAT depot (indicated with red dotted line). **(G)** Fluorescent images of medial and lateral scBAT from *Pax3-mTmG* mice. (green = mG, red = mT, blue = DAPI, Scale = 50 μm) **(H)** Quantification of GFP^+^ cells as a percentage of total adipocytes (n = 3). Data are presented as mean ± SEM.

## RESULTS

### *Pax3*^+^ progenitors do not give rise to supraclavicular brown adipocytes

*Pax3*, together with its orthologue *Pax7*, initiate a transcriptional cascade including *Myf5* and *Myod* for myogenesis. To determine if supraclavicular BAT arises from *Pax3*^+^ myogenic progenitors, we mated *Pax3^Cre^* to *Rosa26^LSL-mT/mG^* reporter mice (**Figure 1C**). *Pax3*-derived cells express membrane-tethered GFP (mG), while those non-*Pax3* progeny cells express membrane-tethered tdTomato (mT). As expected, brown adipocytes within dorsal BAT including interscapular, subscapular and supraspinal depots were almost exclusively GFP^+^ (**Figure 1D, E**), indicating their *Pax3*-lineage identity. Mouse scBAT localizes in the ventral site of the neck, beneath the submandibular gland and tightly connected to the jugular vein (**Figure 1F**). We dissected both medial and lateral scBAT depots, which are above and below the jugular vein respectively. Nearly none of the brown adipocytes in these scBAT depots were GFP^+^ (**Figure 1G, H**). These data demonstrate that scBAT adipocytes are not progeny of somite myogenic progenitors.

### *Myf5*^+^ progenitors do not give rise to scBAT adipocytes

*Myf5*^+^ myogenic progenitors contribute to trunk and limb muscles, and dorsal adipose tissues including iBAT. To determine if scBAT arises from *Myf5*^+^ cells, we mated *Myf5^Cre^*to *Rosa26^LSL-mT/mG^* reporter mice (**Figure 2A**). Similar to the *Pax3^Cre^* reporter, adipocytes within dorsal BAT depots were mostly mG^+^ (**Figure 2B, D**), representing their somite origins. However, only ∼7% of adipocytes in the medial scBAT were GFP-labelled and essentially no adipocytes were labelled in the lateral scBAT (**Figure 2C, D**). Collectively, our *Pax3^Cre^* and *Myf5^Cre^* fate tracing date demonstrate that scBAT and iBAT do not share the same myogenic origins.

**Figure 2.**
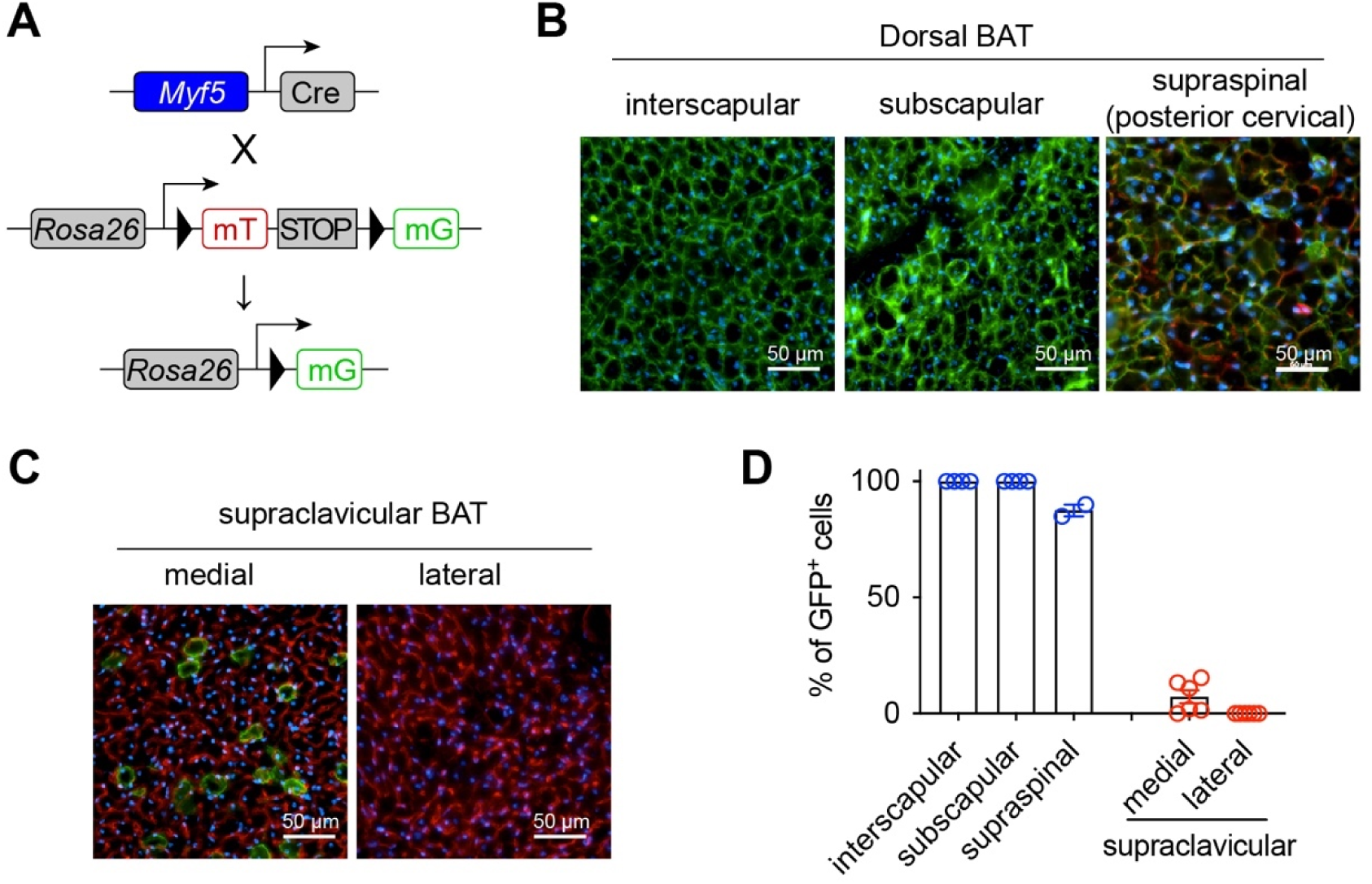
scBAT does not arise from *Myf5*^+^ precursor cells. **(A)** Generation of the reporter mice for the *Myf5*^+^ lineage cells. **(B-C)** Fluorescent images of dorsal BAT (B) or scBAT (C) from 2-month-old female *Myf5-mTmG* mice (green = mG, red = mT, blue = DAPI, Scale = 50 μm). **(D)** Frequency of GFP^+^ cells within each BAT depot (n = 2-5). Data are presented as mean ± SEM.

To validate the lineage tracing data, we then generated *Pparg* knockout mice specifically in *Myf5*^+^ cells (*Pparg^ΔMyf5^*). The *Pparg* gene encodes the master transcriptional factor for adipogenesis - peroxisome proliferator-activated receptor gamma (PPARγ). As a result, severe BAT paucity was observed in the interscapular and subscapular depots (**Figure 3A, B**). Less obvious mass reduction was seen in the supraspinal BAT (**Figure 3A, B**), possibly due to the existence (∼15%) of non-*Myf5*^+^ progeny cells in this depot (**Figure 2B, D**). Western blotting showed a complete loss of PPARγ protein and significant reduction in UCP1 expression (**Figure 3C**). In contrast, scBAT didn’t reduce its size in *Pparg^ΔMyf5^* mice (**Figure 3D**). Instead, there was a trending increase in weight when compared to littermates (**Figure 3E**), suggesting a potential compensation for iBAT paucity. *Myf5^Cre^*does not target scBAT, thus no change in PPARγ or UCP1 expression was observed (**Figure 3F**). Taken together, *Myf5*^+^ myogenic progenitors are essential for the development of dorsal-located BAT, but not ventral-located scBAT.

**Figure 3.**
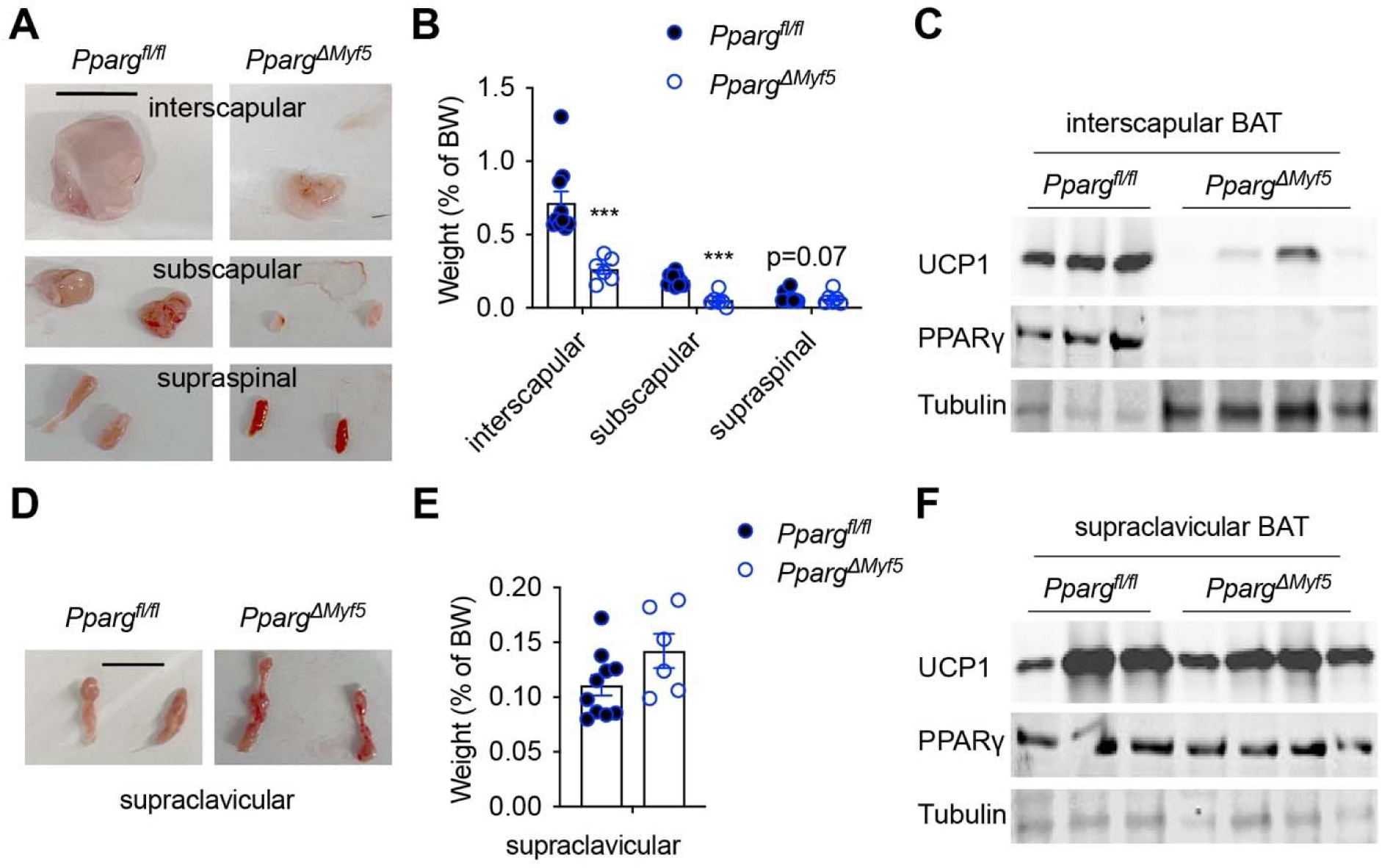
Intact scBAT in mice with PPARγ deficiency in the *Myf5*^+^ lineage. **(A, B)** Indicated dorsal BAT depots from 8-month-old *Pparg^f/f^* (mixed sex, n = 10) and *Pparg^ΔMyf5^* mice (mixed sex, n = 6) were isolated, photographed (A, scale = 1 cm), and weighed (B). **(C)** Immunoblotting of UCP1 and PPARγ with iBAT proteins from *Pparg^f/f^* (n = 3) and *Pparg^ΔMyf5^* (n = 4) mice. **(D, E)** scBAT depots were isolated from 8-month-old *Pparg^f/f^*(n = 10) and *Pparg^ΔMyf5^* mice (n = 6), photographed (D, scale = 1 cm), and weighed (E). **(F)** UCP1 and PPARγ expression in iBAT from *Pparg^f/f^* (n = 3) and *Pparg^ΔMyf5^* (n = 4) mice. Data are presented as mean ± SEM. ***, p < 0.01 by unpaired student’s t-test.

### scBAT adipocytes arise from *Tbx1*^+^ progenitors

The *Tbx1* gene is expressed in cardiopharyngeal mesoderm (CPM) that gives rises to the branchiomeric and transition zone muscles between head and truck. To determine if *Tbx1*^+^ progenitors mark the nearby scBAT, we generated *Tbx1^Cre^*-dependent mT/mG reporter mice (**Figure 4A**). The tracing of the CPM was confirmed by the mT-labeling of scapular muscle (*Myf5*-lineage) and the mG-labeling of clavicular muscle (*Tbx1*-lineage) (**Figure S1**). In consistent with their identity as *Myf5*^+^ progeny, brown adipocytes in dorsal BAT, including interscapular, subscapular, and supraspinal depots, were not labelled at all by mG in *Tbx1-mT/mG* mice (**Figure 4B, D**). In contrast, nearly 50% of scBAT adipocytes are mG^+^ (**Figure 4C, D**). We saw the even distribution of *Tbx1*-lineage adipocytes across the whole scBAT, which is preserved in aged mice (**Figure 4E**). Salivary gland can be found to be connected with the medial end of scBAT. Salivary gland is derived from oral epithelium, thus not labelled by mG (**Figure 4E**). It is currently unclear what the identity of these mT^+^ non-*Tbx1* progeny adipocytes in the scBAT depot.

**Figure 4.**
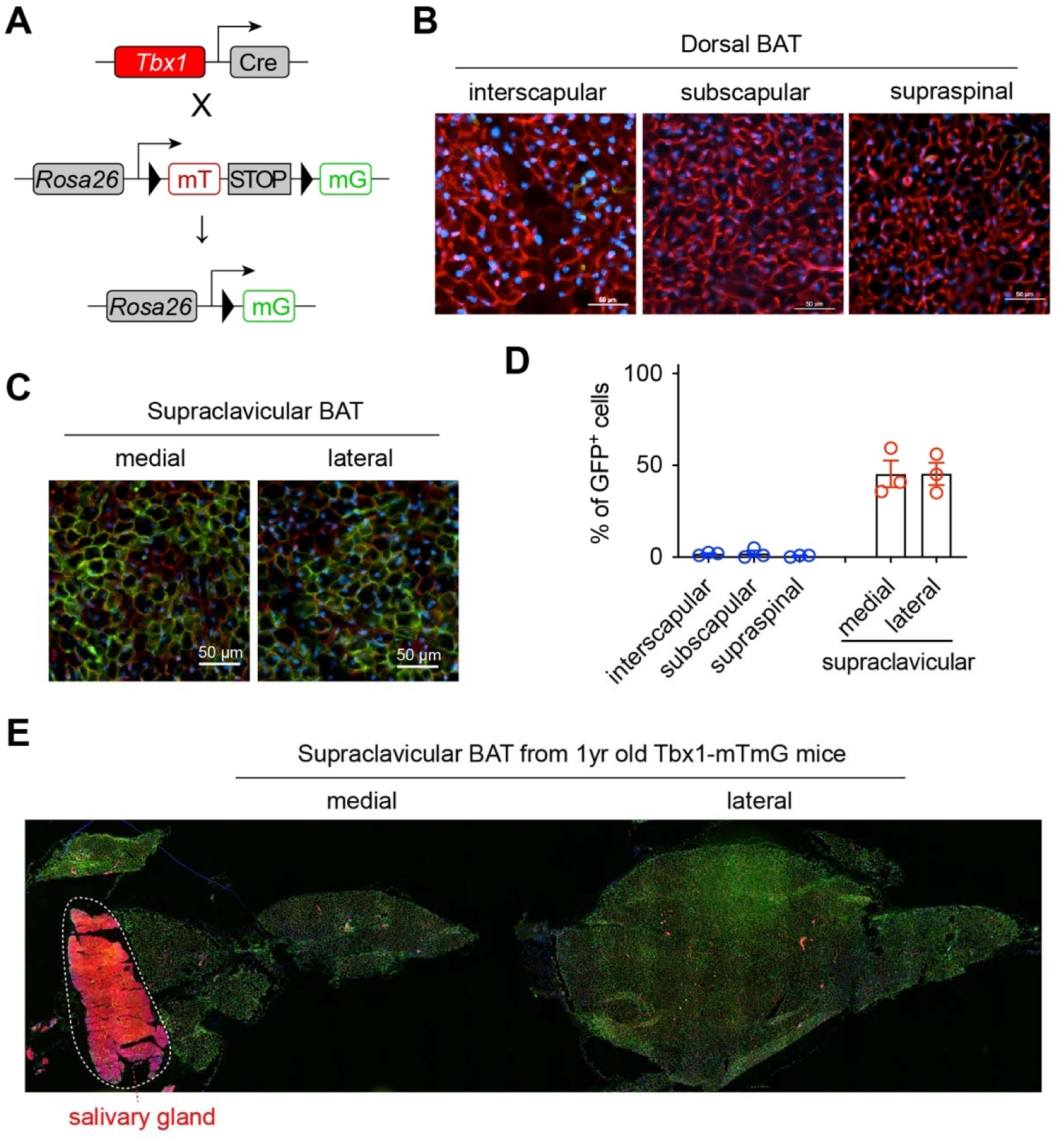
scBAT is composed of *Tbx1*^+^ lineage adipocytes. **(A)** *Tbx1*^+^ lineage cell labelling with the *Tbx1-mTmG* mice. **(B-C)** Fluorescent images of dorsal BAT (B) and scBAT (C) from 6-week-old female *Tbx1-mTmG* mice (green = mG, red = mT, blue = DAPI, Scale = 50 μm). **(D)** Frequency of GFP^+^ cells within each BAT depot (n = 3). Data are presented as mean ± SEM. **(E)** Fluorescent images of scBAT from a 1-year-old *Tbx1-mTmG* mouse (green = mG, red = mT, blue = DAPI, Scale = 50 μm). Salivary gland (epithelial lineage) was labelled by mT.

### scBAT contributes to temperature maintenance in mice

To evaluate the functional contribution of scBAT to systemic metabolism, we generated *Tbx1*-specific *Pparg* knockout mice (*Pparg^ΔTbx1^*). In female *Pparg^ΔTbx1^* mice, PPARγ deficiency leads to a specific decrease of scBAT weight (**Figure 5A**), but not of any dorsal depots including iBAT (**Figure 5B-D**). Body mass and weights of WAT and skeletal muscle were not affected in these *Pparg^ΔTbx1^* female (**Figure S2A, B**). RT-qPCR revealed a significant downregulation of total *Ucp1* transcript in scBAT but not iBAT (**Figure 5E, F**). As a result, female *Pparg^ΔTbx1^* mice reduced more body temperature when challenged with cold, compared to *Pparg^f/f^* controls (**Figure 5G**). Similar scBAT paucity but intact dorsal BAT depots were observed in male *Pparg^ΔTbx1^* mice (**Figure 5H-K**). Body, WAT and muscle weight were comparable between two genotypes in males (**Figure S2C, D**). Histological assessment of scBAT found more unilocular adipocytes in *Pparg^ΔTbx1^*mice (**Figure 5L**), indicative of the loss of brown identity in PPARγ-deficient adipocytes. Nonetheless, male *Pparg^f/f^*and *Pparg^ΔTbx1^* mice had similar tolerance to cold challenge (**Figure 5M**), indicating that male mice rely less on scBAT for thermogenesis.

**Figure 5.**
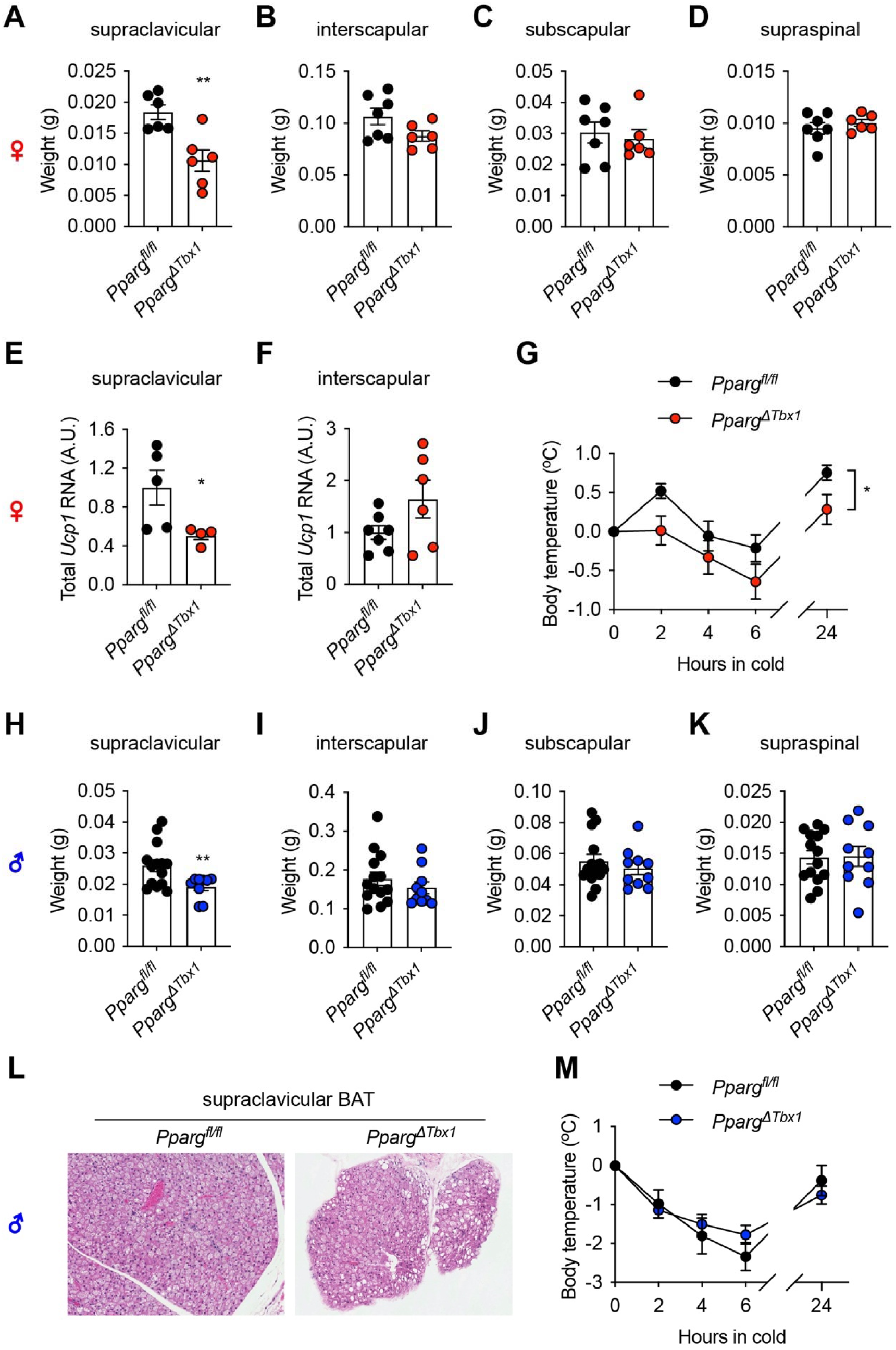
PPARγ deficiency in the *Tbx1*^+^ lineage impairs scBAT function. (**A-D**) Weights of supraclavicular (A), interscapular (B), subscapular (C), and supraspinal (D) BAT depots from 4-month-old *Pparg^f/f^* (n = 7) and *Pparg^ΔTbx1^* (n = 6) female mice. (**E, F**) *Ucp1* gene expression in scBAT (E) and iBAT (F) was determined by RT-qPCR and adjusted by total tissue RNA to calculate the relative total transcript levels. (**G**) Core body temperature of *Pparg^f/f^* (n = 9) and *Pparg^ΔTbx1^* (n = 7) female mice during cold challenge in 4°C. (**H-K**) Weights of supraclavicular (H), interscapular (I), subscapular (J), and supraspinal (K) BAT depots from 4-month-old *Pparg^f/f^* (n = 14) and *Pparg^ΔTbx1^* (n = 10) male mice after 3 weeks of cold challenge. (**L**) Representative H&E staining of scBAT from *Pparg^f/f^* and *Pparg^ΔTbx1^* male mice. (**M**) Core body temperature of *Pparg^f/f^* (n = 6) and *Pparg^ΔTbx1^*(n = 17) male mice during cold challenge in 4°C. Data are presented as mean ± SEM.*, p < 0.05; **, p < 0.01; and ***, p < 0.001 by unpaired student’s t-test or two-way ANOVA (G).

### scBAT paucity does not exacerbate diet-induced obesity

In humans, scBAT prevalence is negatively correlated with BMI and cardiometabolic dysfunction. We thus went on to test if scBAT paucity renders mice more susceptible to high fat diet (HFD)-induced obesity and complications. We subjected both female and male *Pparg^ΔTbx1^* mice and their littermate controls to HFD feeding; however, no difference in weight gain was observed between genotypes (**Figure 6A, B**). *Pparg^ΔTbx1^* mice also had similar tolerance to glucose (**Figure 6C**), suggesting scBAT paucity in mice is not sufficient to cause metabolic dysfunction.

**Figure 6.**
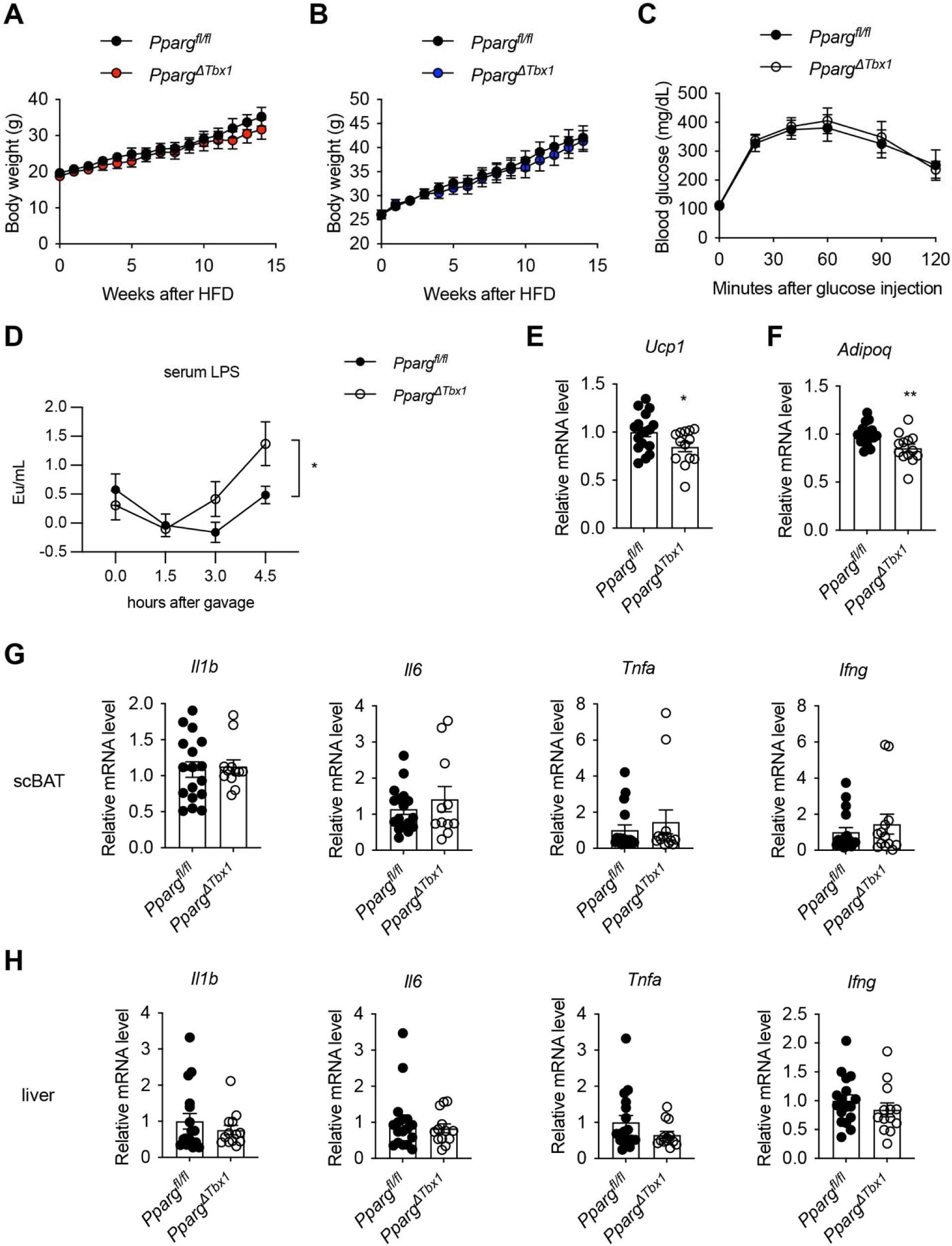
Characterization of HFD-fed *Pparg^ΔTbx1^* mice. (**A, B**) Body weight of female (A, n = 6-7) and male (B, n = 10-12) *Pparg^f/f^*and *Pparg^ΔTbx1^* mice after HFD feeding. (C) Glucose tolerant test of HFD-fed *Pparg^f/f^* (n = 8) and *Pparg^ΔTbx1^* (n = 10) mice (mixed sexes). **(D)** HFD-fed *Pparg^f/f^* (n = 10) and *Pparg^ΔTbx1^* (n = 10) male mice were fasted overnight, orally gavaged with 4 μg LPS (in 200 μl olive oil), and sera were collected at indicated time for LPS measurement. (**E-H**) HFD-fed *Pparg^f/f^* (10 males, 7 females) and *Pparg^ΔTbx1^*(8 males, 5 females) mice were fasted and gavaged with 50 ul of 0.1 μg/ul LPS in water per 20 g of body weight. scBAT (E-G) and liver (H) were collected 3 h later for RT-qPCR quantification the expression of *Ucp1* (E), *Adipoq* (F), and inflammatory genes (G, H). Data are presented as mean ± SEM. *, p < 0.05; **, p < 0.01 by unpaired student’s t-test (E, F) or two-way ANOVA (D).

Obesity induces systemic and hepatic inflammation, and BAT recruitment and UCP1 activation by cold have been suggested to participate in resolving systemic and hepatic inflammation (Mills et al., 2022; Sugimoto et al., 2022). To examine if scBAT is involved in inflammation resolution, we subjected HFD-fed *Pparg^ΔTbx1^*mice to lipopolysaccharide (LPS) injection to generate acute endotoxemia (Munro et al., 2020). Notably, higher serum levels of LPS were observed in *Pparg^ΔTbx1^*mice compared to controls (**Figure 6D**), indicative of possible role of scBAT in neutralizing LPS. As expected, *Ucp1* and *Adipoq* gene expression was downregulated in scBAT of HFD-fed, LPS-treated *Pparg^ΔTbx1^* mice (**Figure 6E, F**). However, we did not observe any significant changes in the expression of inflammatory genes such as *Il1b*, *Il6*, *Tnfa*, and *Ifng* in either scBAT (**Figure 6G**) or liver (**Figure 6H**). We speculate that, because of the intact dorsal BAT depots in these animals, specific paucity of the smaller scBAT would not predispose animals to HFD-induced metabolic dysfunction and inflammation.

### PRMD16 drives the thermogenic programing of scBAT

Next, we investigated the molecular determinants of scBAT development and function. Because of the similar thermogenic activity and regulation between scBAT and iBAT (Mo *et al*., 2017; Shi *et al*., 2021), we postulated that the transcriptional regulatory circuits for iBAT will control scBAT differentiation and/or activity (Kajimura et al., 2010; Shapira and Seale, 2019). PRDM16 dictates the brown adipogenic switch of myogenic progenitors (Seale *et al*., 2008), and is required for WAT browning and the maintenance of brown adipocyte identify (Cohen et al., 2014; Harms et al., 2014). In *Pparg^ΔTbx1^* mice, *Prdm16* gene expression was downregulated in scBAT (**Figure 7A**). To investigate the role of PRDM16 in scBAT, we generated *Tbx1^Cre^*-mediated PRDM16 knockout (*Prdm16^ΔTbx1^*) mice. While we did not observe weight changes in tissues including dorsal BAT depots, scBAT, WAT, and quadricep muscles in female *Prdm16^ΔTbx1^* mice (**Figure 7B**), they showed more body temperature loss during the cold tolerance test (**Figure 7C**). RT-qPCR analysis revealed profound downregulation of thermogenic/adipogenic genes including *Ucp1*, *Pparg*, *Dio2*, *Cidea*, *Pparg1c*, and *Adipoq* specifically in scBAT, but not iBAT of female *Prdm16^ΔTbx1^* mice (**Figure 7D**). Similar findings were observed in male *Prdm16^ΔTbx1^* mice. Compared to littermate controls, male *Prdm16^ΔTbx1^* mice were less cold tolerant (**Figure 7E**), despite of having similar weight of scBAT and other depots (**Figure 7F**). These data suggest that PRDM16 is dispensable for the development of scBAT, but critical for the thermogenic function of scBAT in adult mice.

**Figure 7.**
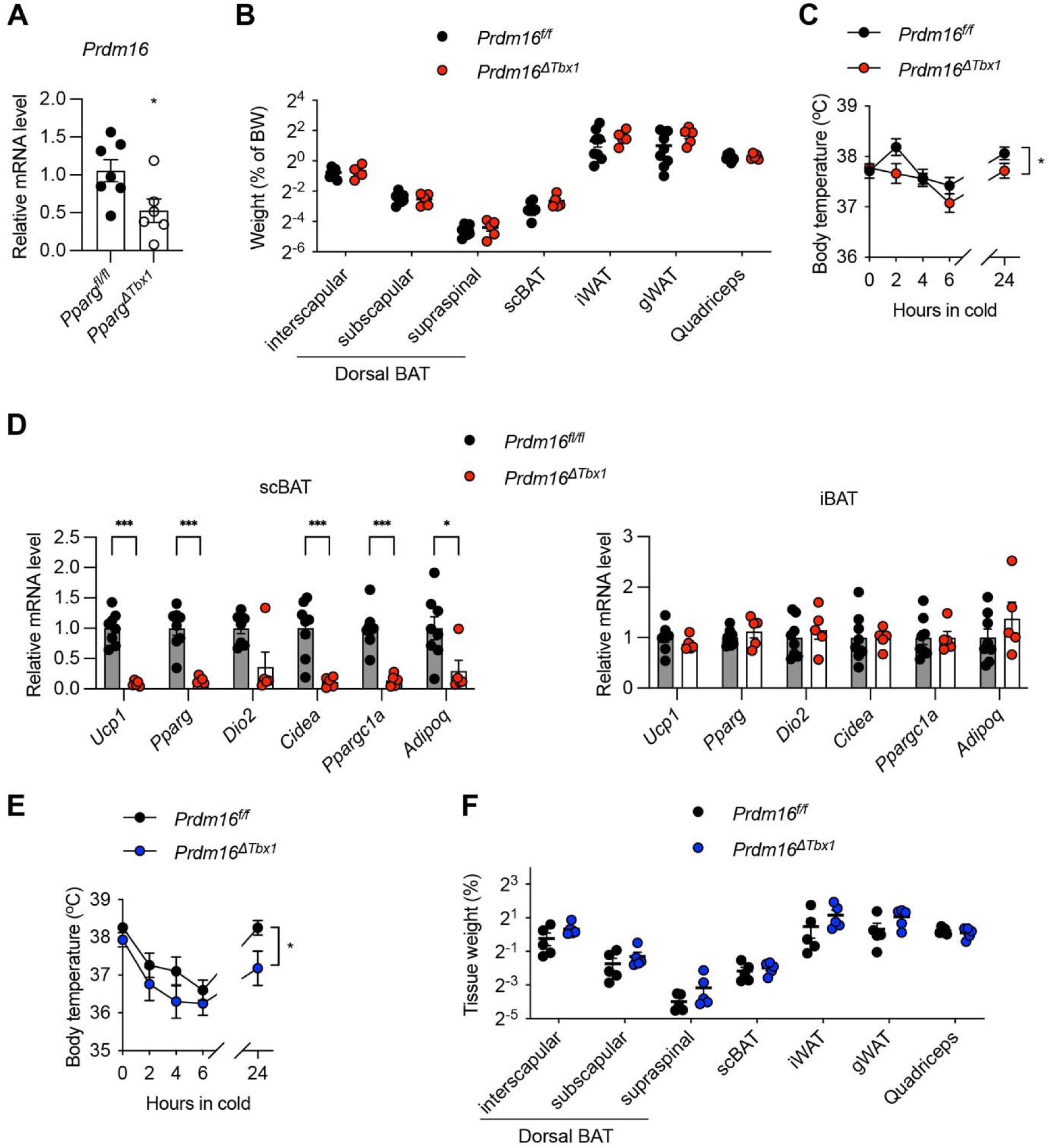
scBAT dysfunction in *Prdm16^ΔTbx1^* mice. **(A)** Expression of *Prdm16* gene in *Pparg^f/f^* (n = 7) and *Pparg^ΔTbx1^* (n = 6) female mice. **(B)** Weight of indicated tissues as a percentage of body weight in *Prdm16^f/f^* (n = 8) and *Prdm16^ΔTbx1^* (n = 5) female mice. **(C)** Core body temperature of *Prdm16^f/f^* (n = 8) and *Prdm16^ΔTbx1^*(n = 5) female mice during cold challenge in 4°C. **(D)** RT-qPCR measurements of gene expression in scBAT and iBAT of *Prdm16^f/f^* (n = 8) and *Prdm16^ΔTbx1^* (n = 5) female mice. **(E)** Core body temperature of *Prdm16^f/f^* (n = 5) and *Prdm16^ΔTbx1^*(n = 6) male mice during cold challenge in 4°C. **(F)** Weight of indicated tissues as a percentage of body weight in *Prdm16^f/f^* (n = 5) and *Prdm16^ΔTbx1^* (n = 6) male mice after 3 weeks of cold challenge. Data are presented as mean ± SEM. *, p < 0.05; **, p < 0.01 by unpaired student’s t-test (A, D) or two-way ANOVA (C, E).

### *TBX1* marks human deep neck BAT

Finally, we determined the expression of *TBX1* gene in human deep neck BAT. Since iBAT was absence in adult humans, subcutaneous neck WAT from healthy donors was obtained as controls (Shaw et al., 2021). RT-qPCR revealed a much higher levels of *TBX1* expression in total lysates from deep neck BAT (**Figure 8A**). Adipogenic differentiation of preadipocytes isolated from both subcutaneous WAT and deep neck BAT induced *TBX1* expression (**Figure 8B**). Nonetheless, a consistent higher expression of *TBX1* could be observed in preadipocytes and adipocytes derived from deep neck BAT, compared to those from subcutaneous neck WAT (**Figure 8B**). Mimicking adrenergic stimulation, the cell-permeable dibutyryl-cAMP stimulates nutrition uptake and oxygen consumption of human brown adipocytes (Arianti et al., 2021). However, cAMP did not change *TBX1* expression (**Figure 8C**), indicating that *TBX1* might not be a cold-inducible early gene that contributes to thermogenesis. Instead, based on our mouse data, *Tbx1* is a location marker specifically for scBAT during development.

**Figure 8.**
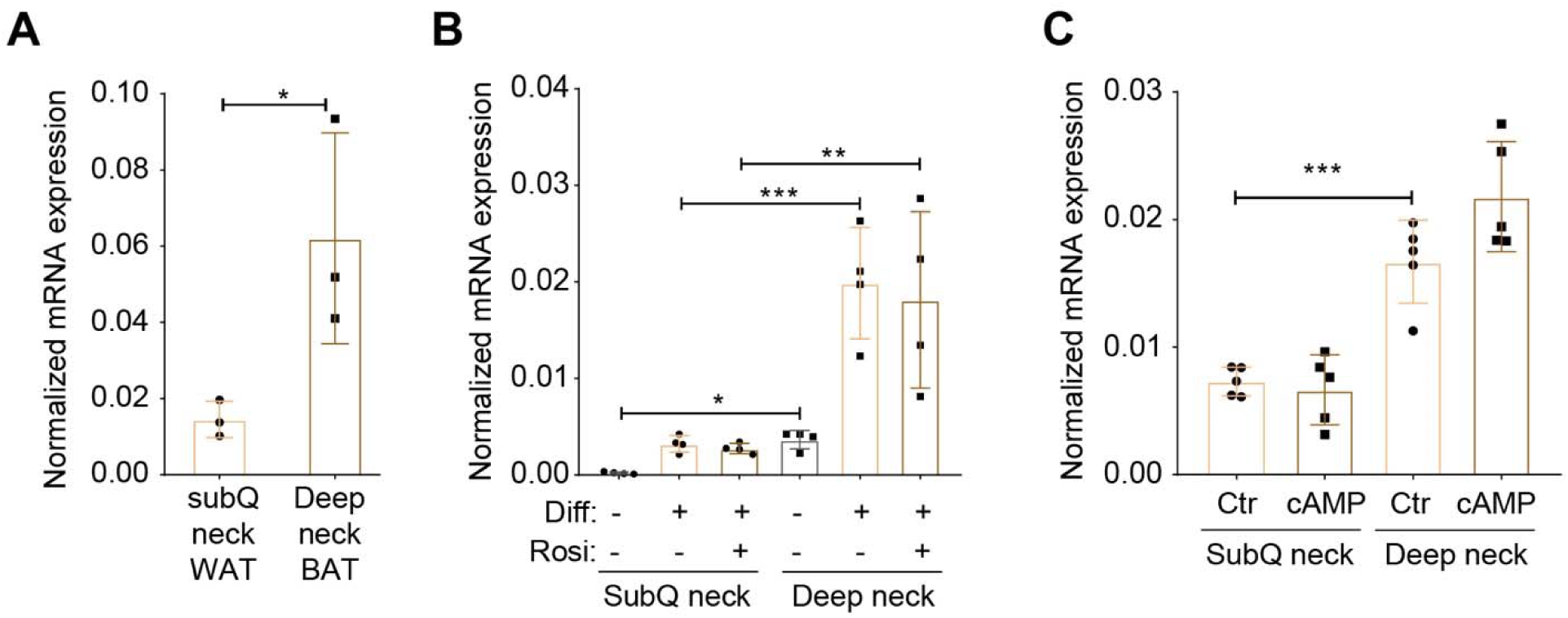
*TBX1* gene expression in human deep neck BAT. **(A)** Taqman RT-qPCR of *TBX1* gene expression, normalized to *GAPDH*, in subcutaneous (subQ) neck WAT and deep neck BAT from 3 donors. **(B)** SubQ neck WAT and deep neck BAT preadipocytes from 4 donors were differentiated to adipocytes in the presence or absence of rosiglitazone for 14 days. *TBX1* gene expression normalized to *GAPDH* was determined using Taqman probes. **(C)** Differentiated adipocytes from SubQ WAT and deep neck BAT (n = 4 donors) were treated with or without 500 μM dibutyril-cAMP for 10 h. *TBX1* gene expression was normalized to *GAPDH*. Data are presented as mean ± SD. *, p < 0.05; **, p < 0.01 by paired student’s t-test (A) or two-way ANOVA (B, C).

## DISCUSSION

Obesity is a major risk factor for many diseases, including type 2 diabetes, cardiovascular disease, and some types of cancers. BAT burns fat and dissipates chemical energy as heat; therefore, activating BAT might be a strategy to combat obesity and related metabolic disorders. Since the re-discovery of supraclavicular BAT (scBAT) in adult humans, many efforts have been devoted in the field to find ways to convert white fat into brown or beige fat. However, almost all the current preclinical studies investigate either interscapular BAT (iBAT) or subcutaneous inguinal WAT in rodents, with assumptions that these depots possess similar developmental, functional, and regulatory mechanisms as the predominant scBAT depot in human adults. In fact, many depot-, location-, and species-specific characteristics of thermogenic adipocytes have been described (Ruan, 2020). For example, human iBAT rapidly atrophies till undetectable in adults (Heaton, 1972; Sidossis and Kajimura, 2015). However, mice iBAT shows little involution or atrophy. Whitened, hypertrophic iBAT adipocytes in aged or “humanized” mice remain as the progeny of *Myf5*^+^ progenitors (Sanchez-Gurmaches and Guertin, 2014), and can be recruited by cold and sympathetic activation (Cui et al., 2016; de Jong et al., 2019; Goncalves et al., 2017; Roh et al., 2018; Sellayah and Sikder, 2014; Tajima et al., 2019). In mice, inguinal WAT adipocytes are derived from *Prrx1*^+^ progenitors in the somatic lateral plate mesoderm (Krueger et al., 2014; Sanchez-Gurmaches et al., 2015). Inguinal WAT browning is driven by *de novo* beige adipogenesis (Berry et al., 2016; Wang et al., 2013), activation of “dormant” beige cells (Rosenwald et al., 2013), and “transdifferentiation” of white adipocytes (Barbatelli et al., 2010; Lee et al., 2015). Like that in mice, subcutaneous WAT in humans can undergoes thermogenic activation (Finlin et al., 2018; Otero-Diaz et al., 2018; Patsouris et al., 2015; Sidossis et al., 2015). However, visceral WAT in humans seems to have higher thermogenic capacity than subcutaneous WAT (Zuriaga et al., 2017).

These depot-and species-specific findings highlight the urgency and importance to understand scBAT development and regulation. Our study here provides the first evidence that scBAT adipocytes do not share the same embryonic origins as iBAT fat cells. Somatic *Pax3*^+^/*Myf5*^+^ myoprogenitors give rise to dorsal-anterior-located iBAT, as well as nearby WAT and trunk muscles. In contrast, *Tbx1*^+^ myoprogenitors from the cardiopharyngeal mesoderm (CPM) give rise to scBAT adipocytes, in addition to many muscle groups in the head and neck. The *Tbx1* gene has been suggested as a marker for beige adipocytes (Wu et al., 2012). However, β3 adrenergic agonism does not induce the expression of *TBX1* in human (Figure 8C) or *Tbx1* in mouse (Ussar et al., 2014). Mouse *Tbx1* was also expressed by inguinal white adipocytes under warm conditions (Roh *et al*., 2018), suggesting that *Tbx1* is not a bona-fide beige marker, but reflects the anatomical location of the inguinal depot. Due to its high expression in inguinal WAT, we speculate, even though not directly examined here, that inguinal fat cells will be labelled in the *Tbx1-mTmG* mice. However, we would not be able to distinguish between their embryonic lineage identity versus postnatal expression of *Tbx1*, because of the constitutive Cre expression in *Tbx1^Cre^* mice. Recently, adipose expression of TBX1 was shown to be necessary for UCP1 expression and insulin sensitivity of subcutaneous WAT (Markan et al., 2020). Future investigations using inducible *Tbx1*-driven Cre models and depot-specific deletion of TBX1 are required to determine the temporospatial function of TBX1. Nonetheless, the shared expression of TBX1 between scBAT and inguinal WAT also provides additional justification to study WAT browning in contributing to metabolic health.

In lineage mapping experiments, we noticed that only about half of scBAT adipocytes are labelled by *Tbx1^Cre^*. This could be a result of low Cre expression or insufficient recombination, supported by the uniform distribution and no cluster formation of mG^+^ adipocytes (Figure 4E). Nonetheless, it is also possible that scBAT adipocytes have multiple developmental origins and *Tbx1* only labels a portion of CPM myoprogenitors. It would be interesting in the future to test whether other CPM markers such as *Ptx2* and *Islet1* (Grimaldi *et al*., 2022; Heude *et al*., 2018; Lescroart *et al*., 2010) trace all or some of the scBAT adipocytes. Although controlling myogenesis at different locations, both *Tbx1* and *Myf5* elicit a transcriptional cascade including *Myod* and *Myogenin* (Braun and Gautel, 2011). *Myod* also induces glycolytic beige adipocytes in the absence of β-adrenergic receptor signaling (Chen et al., 2019). However, *Myod* does not label any iBAT (Sanchez-Gurmaches and Guertin, 2014) or scBAT (our unpublished observations) brown adipocytes. It stresses that the bifurcation between brown adipogenesis and myogenesis happens upstream of *Myod*. The PRDM16-C/EBPβ-EHMT1 transcriptional complex specifies the iBAT brown adipocyte fate from *Myf5*^+^ progenitors and the PRDM16-PPARγ-PGC-1α complex drives the complete brown fat differentiation (Kajimura et al., 2009; Ohno et al., 2013). Here, we have showed that both PPARγ and PRDM16 are also important for scBAT development and function. *Tbx1^Cre^*-mediated deletion of either PPARγ or PRDM16 leads to scBAT dysfunction, evidenced by reduced *Ucp1* expression and cold intolerance. While current evidence suggests shared regulatory mechanisms for the recruitment and activation of iBAT and scBAT (Mo *et al*., 2017; Shi *et al*., 2021), it warrants further investigations to identify depot-specific regulations and functions of BAT, in addition to their distinct developmental origins.

While the absolute mass and thermogenic capacity of human BAT are difficult to quantify, it is well accepted that BAT prevalence declines as a function of age (Huang *et al*., 2022; Ruan, 2020). Thus, many have questioned the physiological relevance of BAT in thermogenesis and body temperature control in adult humans, particularly the elderly. The thermogenic contribution of human BAT might be limited; however, it is without doubt that the presence of scBAT is independently correlated with lower incidences of obesity, type 2 diabetes, dyslipidemia, hypertension, and heart failure (Becher *et al*., 2021). It is thus tempting to hypothesize that iBAT is a critical thermogenic organ for infants, while scBAT in adults primarily modulates systemic metabolism. Anatomically, scBAT sits adjacent to the jugular vein and subclavian vein, where the lymphatic vessels empty lymph collected from the intestine into the venous system. We hypothesized that this unique anatomical location of scBAT may enable its ability to sample and regulate lymphatic fluids, such as bacterial endotoxins therein. While we did see higher serum LPS levels in *Pparg^ΔTbx1^*mice with scBAT paucity (Figure 6D), glucose metabolism and inflammation resolution seemed to be unaffected. The lack of metabolic dysfunction and endotoxemia in *Pparg^ΔTbx1^* mice could be due to the partial atrophy (∼50%) of scBAT and functional compensation from the intact iBAT. Future *Tbx1^Cre^*-mediated brown adipocyte ablation and simultaneous removal of iBAT could address these issues.

In summary, the identification of *Tbx1^+^* lineage cells as progenitors of scBAT brown adipocytes reveals location-specific myoprogenitors for different BAT depots in rodents and possibly humans. This knowledge can be leveraged to develop new models in order to discover depot-specific BAT functions. Not only should we cease to state that “brown adipocytes are derived from a *Myf5*-expressing lineage”, but also study more the *Tbx1^+^* lineage-derived brown adipocytes due to their human relevance.

## METHODS

### Animals

All animal experiments were approved by the institutional animal care and use committee of the University of Minnesota. All the mice were group-housed in light/dark cycle-(6am-8pm light), temperature-(21.5 ± 1.5 °C), and humidity-controlled (30-70%) room, and had free access to water and regular chow (Teklad #2018), or 60% high fat diet (Research Diet #D12492) as indicated. *Tbx1^Cre^* (MGI:3757964) was a kind gift from Dr. Antonio Baldini (Xu et al., 2004). *Myf5^Cre^* (Jax #007893), *Pax3^Cre^*(Jax #005549). *Pparg^f/f^* (Jax #004584), *Rosa26^LSL-mT/mG^*(Jax #007676) mice were from Jackson Lab.

For cold treatment, mice were housed in a temperature-controlled room (4°C) with free access to water and food. Core body temperature was measured using an electronic thermometer with anal probe.

For glucose tolerance test, mice were fasted overnight for 16h, and intraperitoneally injected with 1.5 g/kg body weight of glucose. Blood glucose levels were measured using a Bayer Contour Glucometer at indicated time point after injection.

For LPS treatment, mice were fasted overnight when indicated, and orally gavaged with LPS from Escherichia coli O111:B4 (Sigma, L3024) in water or olive oil. Tail blood was collected for LPS measurement using a Pierce chromogenic endotoxin quantification kit (ThermoFisher, A39552).

### Human adipocytes

Tissue collection was approved by the Medical Research Council of Hungary (20571-2/2017/EKU) followed by the EU Member States’ Directive 2004/23/EC on presumed consent practice for tissue collection. All experiments were carried out in accordance with the guidelines of the Helsinki Declaration. Written informed consent was obtained from all participants before the surgical procedure. During thyroid surgeries, a pair of deep neck BAT and subcutaneous WAT samples was obtained to rule out inter-individual variations. Patients with known diabetes, malignant tumor or with abnormal thyroid hormone levels at the time of surgery were excluded. Tissue specimens were either homogenized in Trizol or digested in phosphate buffered saline (PBS) with 120 U/ml collagenase (Sigma, C1639) to obtain stromal-vascular fraction (SVF). Floating cells were washed away with PBS after three days of isolation and the remaining cells were cultured (Shaw *et al*., 2021). Human primary adipocytes were differentiated from SVF of adipose tissue containing preadipocytes according either to a regular adipogenic protocol or in the presence of long-term rosiglitazone effect resulting in higher browning capacity of the adipocytes. Where indicated, adipocytes were treated with a single bolus of 500 µM dibutyryl-cAMP (Sigma, D0627) for 10 hours to mimic *in vivo* cold-induced thermogenesis (Arianti *et al*., 2021). Then, SVF cells or adipocytes were homogenized using Trizol.

### Histology

Brown adipose tissues were harvested and fixed in 10% formalin overnight with gentle shaking, then kept in 4°C for further experiments. For hematoxylin and eosin staining, BAT embedding, sectioning, staining was conducted at the Comparative Pathology Shared Resource of the University of Minnesota. For the fluorescent imaging, the fixed BAT was embedding with OCT (Tissue-Tek #4583) then sliced into 10 μm slides. Following PBS washing, the sections were mounted using VECTASHIELD® Antifade Mounting Medium with DAPI and visualized using Nikon Ni-E or Keyence microscope system. The numbers of Tomato^+^ and GFP^+^ cells were counted to calculate the percentage of GFP^+^ cells.

### RT-qPCR

After weight measurement, BAT tissues were homogenized in Trizol (Thermo Scientific) for RNA isolation, following the manufacturer’s protocol. RNA concentrations were measured with a NanoDrop spectrophotometer. Reverse transcription was performed with the iScript™ cDNA Synthesis Kit. Real-time RT-PCR was conducted using iTaq™ Universal SYBR® Green Supermix and gene-specific primers on a Bio-Rad C1000 Thermal Cycler. Relative expression was normalized to the house keeping *Rplp0* gene. Total *Ucp1* mRNA amounts were calculated based on relative *Ucp1* mRNA levels and total amounts of RNA isolated from specific depots (Nedergaard and Cannon, 2013). For human WAT, BAT, and adipocyte lysates, RT-PCR was conducted using validated TaqMan assays (ThermoFisher, Hs00271949_m1 for *TBX1*, and Hs99999905_m1 for *GAPDH*). Gene expression values were calculated by the comparative threshold cycle (Ct) method. ΔCt represents the Ct of the target minus that of *GAPDH*. Normalized gene expression levels equal 2−ΔCt (Arianti *et al*., 2021).

### Western blot

Tissues were collected quickly after sacrificing mice and homogenized immediately in RIPA lysis buffer (50 mM Tris–HCl pH 7.4, 1% Nonidet P-40, 0.25% Na-deoxycholate, 150 mM NaCl, 1 mM EDTA and protease inhibitors) on ice-water bath. Protein concentration was measured using a BCA protein assay kit (ThermoFisher). Protein samples were separated by SDS-PAGE and Western blots were performed with following antibodies: UCP1 (Abcam, #ab209483), PPARγ (Cell Signaling Techonology, #2443), Tubulin (Santa cruz, #SC-8035).

### Quantification and statistical analysis

Results are shown as mean ± SEM or ± SD. N values (biological replicates) and statistical analysis methods are described in figure legends. The statistical comparisons were carried out using two-tailed Student’s t-test and one-way or two-way ANOVA with indicated post hoc tests with Prism (Graphpad). Differences were considered significant when p < 0.05. *, p < 0.05; **, p < 0.01; ***, p < 0.001.

## ACKNOWLEGEMENT

We thank Dr. Antonio Baldni for kindly providing the *Tbx1^Cre^*mice and Dr. Bernice Morrow for shipping the mice to us. We thank Dr. Jun Wu and Dr. Rita Perlingeiro for sharing the *Prdm16^f/f^* and *Pax3^Cre^*mice, respectively. We thank Dr. Ferenc Győry for the surgical removal of human BAT and WAT biopsies. This work was supported by the National Natural Science Foundation of China, China (32170847) to Z.H., National Research, Development and Innovation Office of Hungary (NKFIH-FK131424) to E.K., and Department of Integrative Biology and Physiology Grant Accelerator Program to H.-B.R.

## AUTHOR CONTRIBUTION

Z.H. initiated the project, generated mouse lines, performed fate mapping, and contributed to manuscript writing. C.G. characterized *Pparg^ΔTbx1^*and *Prdm16^ΔTbx1^* mice. Z.Z assisted with mouse colony management, fate mapping, and GTT. A.S. quantified frequencies of mG^+^ adipocytes. K.T. performed Western blotting. A.B. assisted with tissue dissection. R.A and E.K. collected human samples and performed RT-qPCR of human cells. H.-B.R. conceived the project, designed experiments, analyzed data, and wrote the manuscript.

---

**Figure S1.**
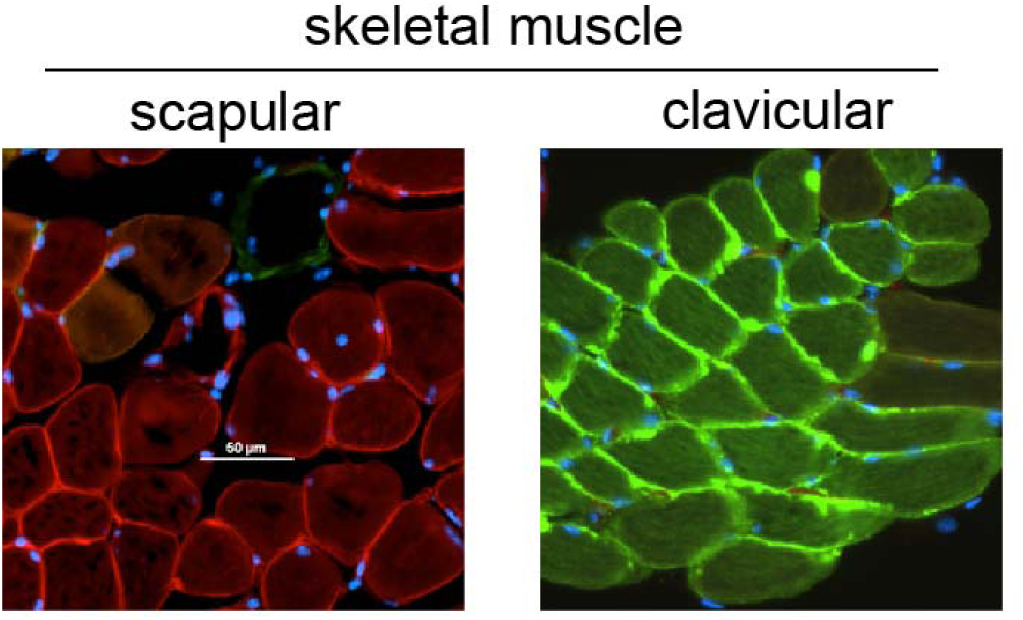
Validation of *Tbx1-mTmG* reporter mice. Representative fluorescent images of scapular (left) and clavicular (right) skeletal muscles from *Tbx1-mTmG* reporter mice (scale = 50 μm).

**Figure S2.**
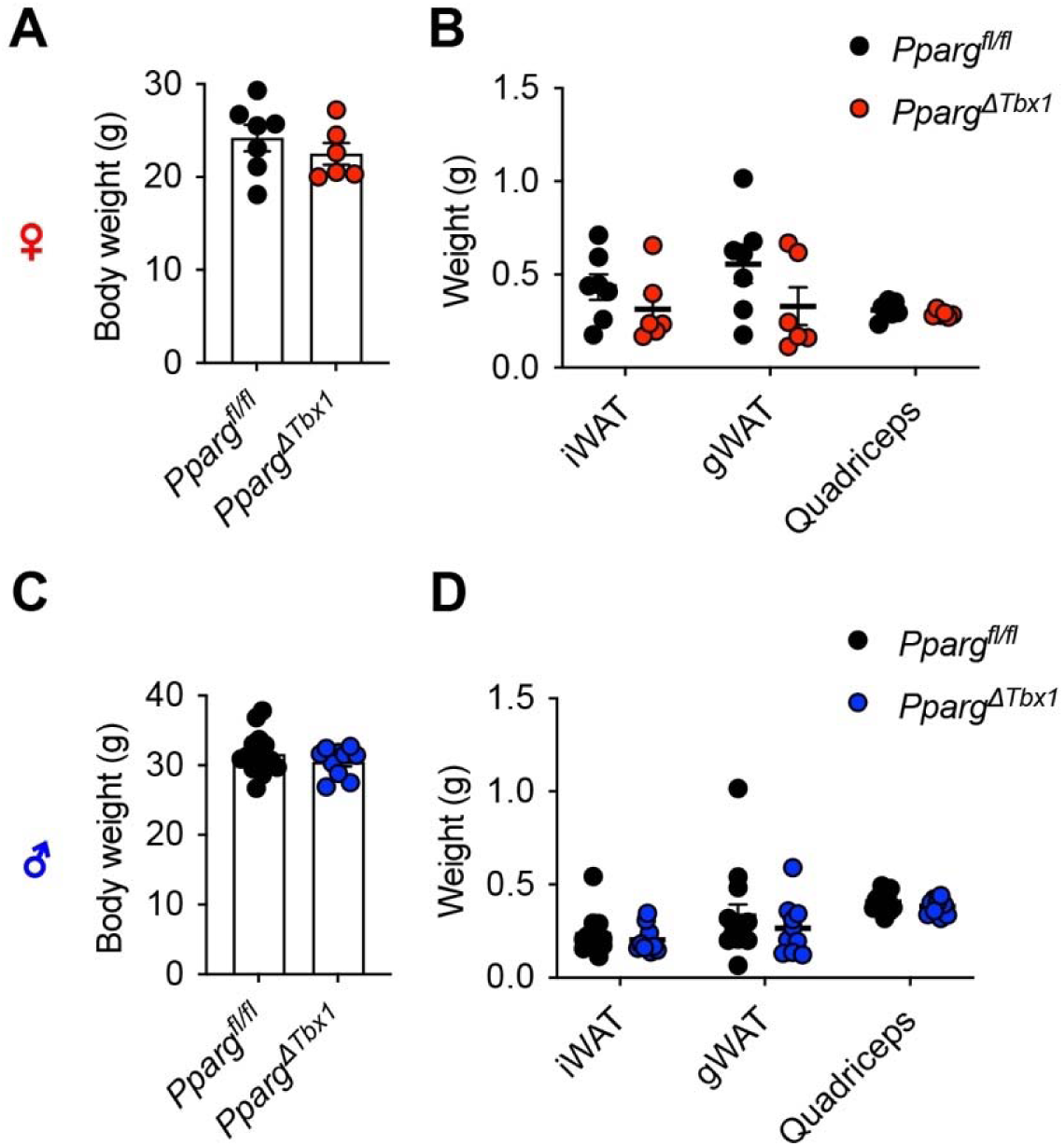
No changes in body and tissue weight of *Pparg^ΔTbx1^* mice. (**A, B**) Body weight (A) and tissue weight (B) of 4-month-old *Pparg^f/f^* (n = 7) and *Pparg^ΔTbx1^* (n = 6) female mice. (**C, D**) Body weight (C) and tissue weight (D) of 4-month-old *Pparg^f/f^* (n = 14) and *Pparg^ΔTbx1^* (n = 10) male mice.

